# Genome-wide analysis of chalcone synthase (CHS) family from eggplant (*Solanum melongena* L.) in flavonoid biosynthetic pathway and expression pattern in response to heat stress

**DOI:** 10.1101/863696

**Authors:** Xuexia Wu, Shengmei Zhang, Xiaohui Liu, Jing Shang, Aidong Zhang, Zongwen Zhu, Dingshi Zha

**Author notes:** Corresponding author: Dingshi Zha. These authors contributed equally to this work (Xuexia Wu, Shengmei Zhang).

## Abstract

Enzymes of chalcone synthase (CHS) family participate in the synthesis of a series of secondary metabolites in plants, fungi and bacteria. CHS showed significant correlation to the accumulation patterns of anthocyanin. The peel color mainly determined by the content of anthocyanin, is a majority economic trait for eggplant affected by heat stress. A total of 7 *CHS*(*SmCHS1-7*) putative genes were identified in genome-wide of eggplant (*S. melongena* L.). The *SmCHS* genes distribute on 7 scaffolds and were classified into 3 clusters. Phylogenetic relationships analysis showed that 73 *CHS* genes from 7 Solanaceae species were classified into 10 groups. *SmCHS5*, *SmCHS6* and *SmCHS7* were continuously down-regulated under 38°C and 45°C treatment, while *SmCHS4* was up-regulated under 38°C but little change at 45°C in peel. Expression profiles of anthocyanin biosynthesis key genes families showed that the PAL, 4CL and AN11 genes were mainly expressed in all five tissues. CHI, F3H, F3’5’H, DFR, 3GT and bHLH1 genes were expressed in flower and peel. Under heat stress, 52 key genes expression level were reduced under heat stress. By contrast, expression patterns of eight key genes similar to *SmCHS4* up-regulated at the 38°C-3h. Comparative analysis of putative CHS protein biochemical characteristics, *cis*-regulatory elements, regulatory network revealed that *SmCHS* genes family have conservation gene structure and functional diversification. *SmCHS* showed two or more expression patterns and execute multiple functions to regulate anthocyanin content. Combined with regulatory networks, it is possible to further understand the regulation mechanism of peel color in eggplant.

## Introduction

Enzymes of chalcone synthase (CHS) is the member of plants-specific type III ployketide synthase (PKS) [1, 2], catalyses the first committed step of the branch of the phenylpropanoid pathway which leads to the synthesis of flavonoids [3, 4]. Flavonoids are well known as a group of plant secondary metabolites that comprise several different classes of compounds such as chalcones, flavones, flavonols isoflavones and anthocyanins. They have a wide variety of biological functions, such as flower pigmentation, protection against UV radiation, pathogen defense, auxin transport and pollen fertility [5-7]. CHS also showed significant correlation to the accumulation patterns of anthocyanin. After heat treatment, Transcript levels of *CHS* decreased in rose flower and in eggplant [8, 9].

The product of the CHS reaction is a pivotal precursor for a vast array of secondary metabolites derived from malonyl-CoA and p-coumaroyl-CoA. CHS exists as homodimeric iterative PKS (monomer size of 42–45 kDa) with two independent active sites that catalyze a series of decarboxylation, condensation, and cyclization reactions [2, 10]. CHS superfamily share high similarity in their amino acid sequence, which contains the structurally conserved catalytic center consisted of four residues Cys-His-Asn-Phe, and most of the genes contain two exons and one intron [11]. But, the *CHS* genes family has been not characterized yet in eggplant.

Anthocyanins are plant secondary metabolites and one of the most abundant natural pigments, that are responsible for the characteristic many colors in flowers, fruits and vegetables plant tissues. The anthocyanin biosynthesis pathway has been studied in numerous plant species and most of the genes involved in this process have been identified. The enzyme evolved in anthocyanin biosynthesis are as follows: phenylalanine ammonia lyase (PAL), cinnamate 4-hydroxylase (C4H), 4-coumarateCoA ligase(4CL), chalcone synthase (CHS), chalcone isomerase (CHI), flavanone 3-hydroxylase (F3H), flavonoid 3′-hydroxylase (F3′H), flavonoid 3′5′-hydroxylase (F3′5′H), dihydroflavonol4-reductase (DFR), anthocyanidin synthase (ANS), and anthocyanidin 3-O-glucosyltransferase (3GT). Besides, Most transcription factors, such as myeloblastosis (MYB) and basic helix-loop-helix (bHLH), are positive regulators of anthocyanin biosynthesis in vegetative tissues [12-14]. Production of chalcone requires the condensation of one molecule of p-coumaroyl-CoA and three malonyl-CoA molecules which is catalyzed by CHS. In conclusion, CHS as the gatekeeper of anthocyanin pathway [15].

Anthocyanins play an important role in plant survival under stressful environmental conditions. high temperatures are known to reduce anthocyanin accumulation and have discoloration effect in many plant tissues. even drastically in colored flowers [8, 16], and the skin of fruits such as grape berries, apples and eggplant [9, 17-20].

Eggplant (S. *melongena* L.) is one of the most important thermophilic vegetable which is produced in many tropical and temperate regions of the world. The optimum growth temperature for eggplant is between 22 and 30 °C. Eggplants subjected to high temperature may lead to stagnation of growth, abortion of flower buds, decrease of pollen viability rate and fruit set, and the peel’s color will turn light when the temperature is over 35 °C. High temperature severely reduce the yield and affects the appearance quality of eggplant. However, the molecular mechanism of high temperature stress in eggplant, are poorly understood.

In current study, all *SmCHS* family members were identified in eggplant. The comprehensive survey of members was performed including gene structures, The biochemical characteristics of putative CHS protein, promoters *cis*-elements, phylogenetic relationships among members in other relative species, as well as their expression profiles in various organs/tissues under high temperature stress. The findings of the present study will provide a new start for functional studies on eggplant *SmCHS* family genes.

## Materials and methods

### Plant materials and RNA extraction

Eggplant cultivar ‘Tewangda’ is a cold tolerance cultivar with blackish purple skin. It is grow vigorous and have good fruit setting. The fruit shape is 27.6cm fruit length, 5.4cm transverse diameter and 209g single fruit weight on average. The ‘Tewangda’ fruit has good commercial property and good transportation resistance. ‘Tewangda’ were grown at the same growth stage were randomly selected. These plants were grown 144 days after sowing, then put inside incubators set at 27°C (CK), 38°C or 45°C for 3 or 6 hours (three plants per treatment). For each treatment, the tissue samples of root, stem, leaf, flower and peel were obtained and immediately frozen in liquid nitrogen, and stored at −80°C for RNA extraction and other analyses. All plant materials examined in this study were obtained from Shanghai Academy of Agricultural Sciences. The total RNA was extracted from each tissue sample by mirVana miRNA Isolation Kit (Ambion) following the manufacturer’s protocol. The extracted total RNA was stored at −80°C. RNA integrity was evaluated using the Agilent 2100 Bioanalyzer (Agilent Technologies, Santa Clara, CA, USA).

### Identification of the CHS family members in eggplant genome

The whole protein sequence of Solanum melongena L. (eggplant) were obtained from the Eggplant Genome DataBase (http://eggplant.kazusa.or.jp) [21], and those of Solanum tuberosum L. (potato, http://solanaceae.plantbiology.msu.edu/pgsc_download.shtml) [22], Solanum lycopersicum (tomato, https://solgenomics.net/organism/Solanum_lycopersicum/genome) [23], Solanum penellii (wild tomato, https://www.plabipd.de/project_spenn/start.ep) [24], Capsicum annuum L. (pepper, http://peppergenome.snu.ac.kr) [25], Petunia axillaris (https://solgenomics.net/organism/Petunia_axillaris/genome) [26], Petunia inflate (https://solgenomics.net/organism/Petunia_inflata/genome) [26], Nicotiana tabacum (common tobacco, https://www.ncbi.nlm.nih.gov/nuccore/AYMY00000000) [27]. The profiles of CHS (PF00195 and PF02797) was download from Pfam protein family database (http://pfam.xfam.org/) and these profile sequence were used as query to perform BLASTP search against the protein sequence data of all the species which mentioned above with a maximum E-value of 1×10^−3^, respectively [28]. To further verify the exact copy number of CHS and remove redundant sequences, the Pfam database and Genome websites were also searched using “chalcone synthase” as keyword. All CHS sequences were submitted to EXPASy (https://web.expasy.org/protparam/) to calculate the number of amino acid, molecular weight and theoretical isoelectric point (pI).

### Structure characterization

The locations and intron numbers of CHS were acquired through the genome website. All of the acquired protein sequences wreath first aligned by ClustalX software with the default parameters [29]. An unrooted maximum-likelihood phylogenetic tree was constructed using MEGA6 software with bootstrap test of 1000 times [30]. The MEME program (Version 5.0.5, http://meme-suite.org/tools/meme) was used to identified the conserved motif of the CHS sequences with the following parameters, any number of repetitions, maximum of 10 misfits and optimum motif width of 6-200 amino acid residues. The WoLF PSORT program was used to predict the subcellular localization information of CHS proteins (https://www.genscript.com/wolf-psort.html) [31].

### Analysis of cis–acting element in SmCHS

The upstream sequences (2 kb) of the *SmCHS* coding sequences in eggplant were retrieved from the genome sequence, and then submitted to PlantCARE (http://bioinformatics.psb.ugent.be/webtools/plantcare/html/) to identify regulatory elements [32].

### Phylogenetic analysis of CHS genes

The full-length protein sequences of all eight species in Solanaceae were used for phylogenetic analysis. All of the protein sequences were first aligned by ClustalX software with the default parameters [29]. The phylogenetic tree was generated with MEGA6 software with bootstrap test of 1000 times. The final tree was viewed and modified in Evolview software [33]. The CHS genes were classified into different groups according to the topology of phylogenetic tree.

### Construction of the mRNA regulatory network

RNA-seq results come from our lab [34]. The significant differentially expressed genes (Fold change ≥ 2 and *p* value < 0.05) were used to calculate pearson correlation coefficient between *CHS* genes and others. The TBtools program was used to elucidate the Gene Ontology (GO) functional classification for the mRNAs which with the correlation coefficient more than 0.9 [35]. The top 5 regulatory mRNAs which annotated by GO enrichment for the genes associated with anthocyanin biosynthesis were collected to construct the regulatory network. The network was visualized using Cytoscape [36].

### qRT-PCR analysis

The first-strand cDNA was synthetised from 1 μg of 5 tissues (root, stem, leaf, flower and peel) total RNA using Prime Script RT Reagent Kit (Takara, Dalian, China). The qRT-PCR reactions were performedin 96-well plates using the ABI 7500 fast Real-Time PCR system (Applied Biosystems, USA) with the QuantiFast SYBR Green PCR Kit (Qiagen, Duesseldorf, Germany). The qRT-PCR parameters were as follows: 95 °C for 5 min, then 45 cycles of 95 °C for 10 s, 60 °C for 10 s, and 72°C for 10 s. The relative mRNA expression levels were calculated using the 2^−ΔΔCT^ method [37]. PGK(JX154676) was used as an internal control to normalize the data. The primer sequences are listed in Additional file3 Table S1.

## Results

### Identification of CHS genes and sequence analysis in Solanaceae species

A total of 7 *CHS* (*SmCHS1-7*) genes in eggplant were identified after verified by protein sequences analysis and Blast search by using eggplant genome annotation database (Additional file1 Table S1a). The length of SmCHS protein ranged from 327 to 396 amino acids (Table 1). The molecular weights of SmCHS were between 35.2 kDa and 43.7 kDa. The theoretical pI value of SmCHS ranged from 5.59 to 7.04. In addition, 66 *CHS* genes were characterized from other 7 Solanaceae species. The subfamily numbers of *CHS* genes were ranged from 6 (*Solanum penellii*) to 13 (*Petunia axillaris*) (Table 1, Additional file1 Table S1b-h). The molecular weights of CHS for other 7 Solanaceae species range from 17.3 to 47.4, length of protein ranged from 156 to 431 amino acids, the theoretical pI value ranged from to 5.1 to 8.47 (Additional file2 Table S1a-g). The average of number of amino acid, molecular weight and theoretical pI were calculated, and then as a data set for each specie. The correlation coefficient between the above data were all more than 0.99. This suggests that *CHS* genes are conservative in Solanaceae species.

**Table 1.**
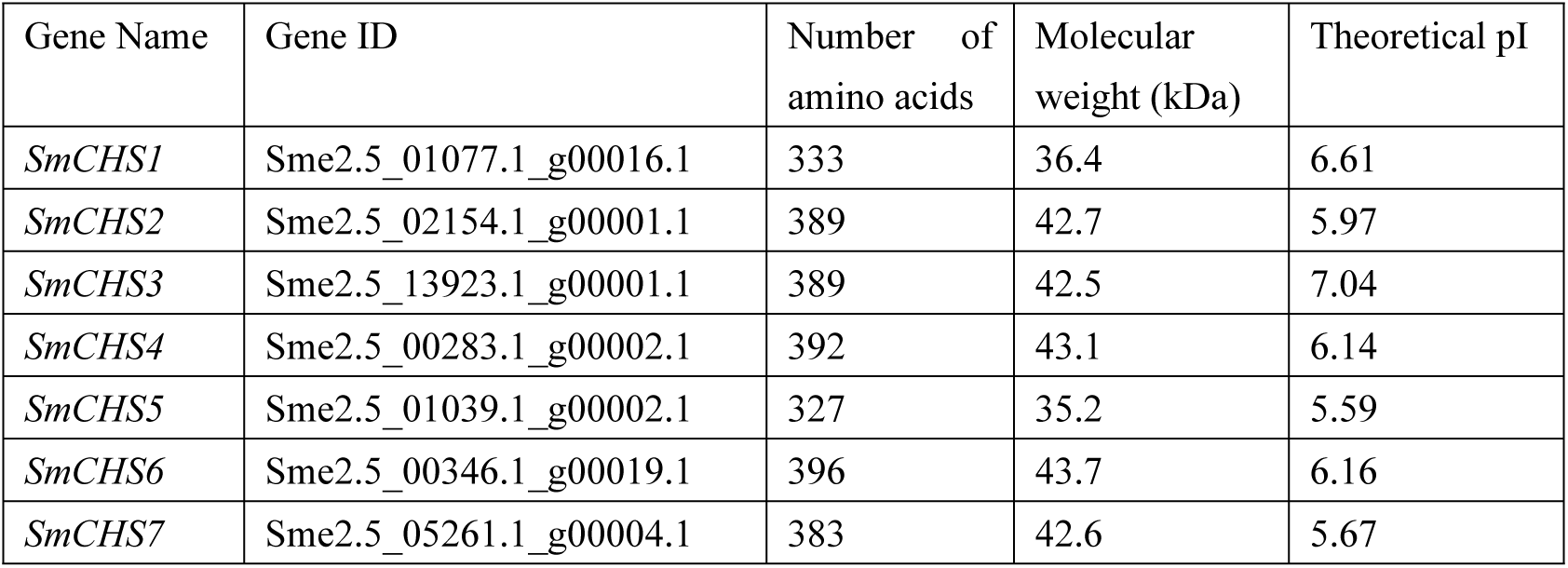
Feature of *SmCHS* genes identified in eggplant.

| Gene Name | Gene ID | Number of amino acids | Molecular weight (kDa) | Theoretical pI |
| --- | --- | --- | --- | --- |
| *SmCHS1* | Sme2.5_01077.1_g00016.1 | 333 | 36.4 | 6.61 |
| *SmCHS2* | Sme2.5_02154.1_g00001.1 | 389 | 42.7 | 5.97 |
| *SmCHS3* | Sme2.5_13923.1_g00001.1 | 389 | 42.5 | 7.04 |
| *SmCHS4* | Sme2.5_00283.1_g00002.1 | 392 | 43.1 | 6.14 |
| *SmCHS5* | Sme2.5_01039.1_g00002.1 | 327 | 35.2 | 5.59 |
| *SmCHS6* | Sme2.5_00346.1_g00019.1 | 396 | 43.7 | 6.16 |
| *SmCHS7* | Sme2.5_05261.1_g00004.1 | 383 | 42.6 | 5.67 |

### Structure and conserved motif analysis of *SmCHS*

The 7 *SmCHS* genes distribute on 7 scaffolds. To better understand the evolution of *SmCHS* genes, an unrooted maximum-likelihood tree was constructed based on the 7 SmCHS protein sequences, and the *SmCHS* were classified into 3 clusters (i, ii and iii) (Fig 1). Among the *SmCHS* genes, only one *SmCHS7* had three exons, the others had two exons (Fig 1) based on available information from the genome annotation. These results suggest the potential biological function diversities of the *SmCHS* genes in eggplant.

**Fig 1.**
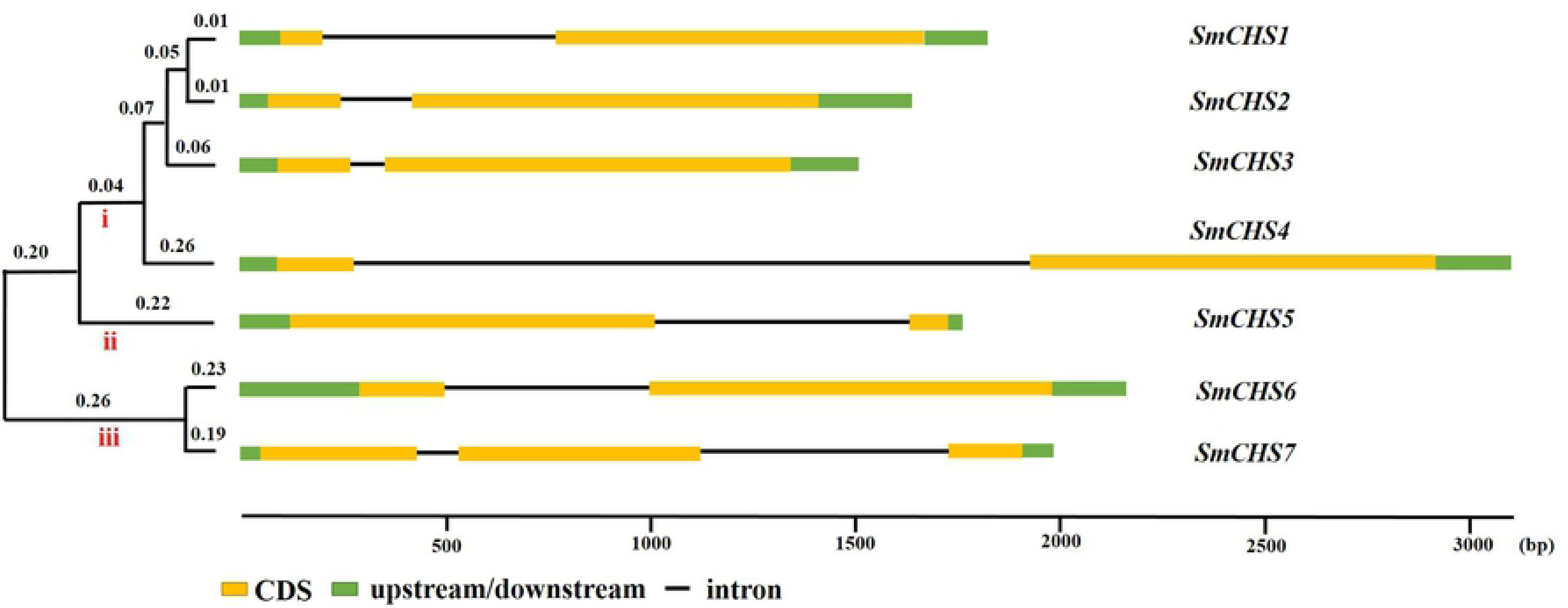
Phylogenetic relationship and gene structure analysis of *SmCHS* genes.

The phylogenetic tree was on the left of the figure, which showed that *SmCHS* were classified into three clusters (i, ii and iii). The exon/intron organization of *SmCHS* was on the right of the figure. For *SmCHS* genes organization, yellow boxes represent exons, black lines represent the intron, and green boxes are indicated in. upstream/downstream region. The length of exon and intron are drawn to scale.

To understand the function diversification of *SmCHS*, the conserved motifs of these 7 protein sequences were identified by MEME program, 10 conserved motifs were detected in eggplant (Fig 2, Table 2). For all the 7 eggplant SmCHS proteins, Motif 1 and Motif 2 exist in all of them, Motif 3 is only absent in *SmCHS5*, Motif 4 and Motif 5 are only absent in *SmCHS1*. The N-terminal domain (PF00195) of CHS protein contained Motif 1 and the combination of Motif 3, 4, 6, 7 and 9. The C-terminal domain (PF02797) of CHS protein contained Motif 2 and the combination of Motif 5, 8 and 10. Therefore, the motif configuration of the SmCHS reflects the conservation and diversity of the CHS family. To further investigate subcellular localization information of SmCHS proteins, WoLF PSORT program was used to predict the localization of SmCHS protein [31]. SmCHS7 was predicted to localize in nuclear, SmCHS4 and SmCHS6 were predicted to localize in chloroplast. The others SmCHS proteins were predicted to localize in cytoplasmic. The different compositions of the domains and subcellular localization may indicate functional diversity.

**Table 2.**
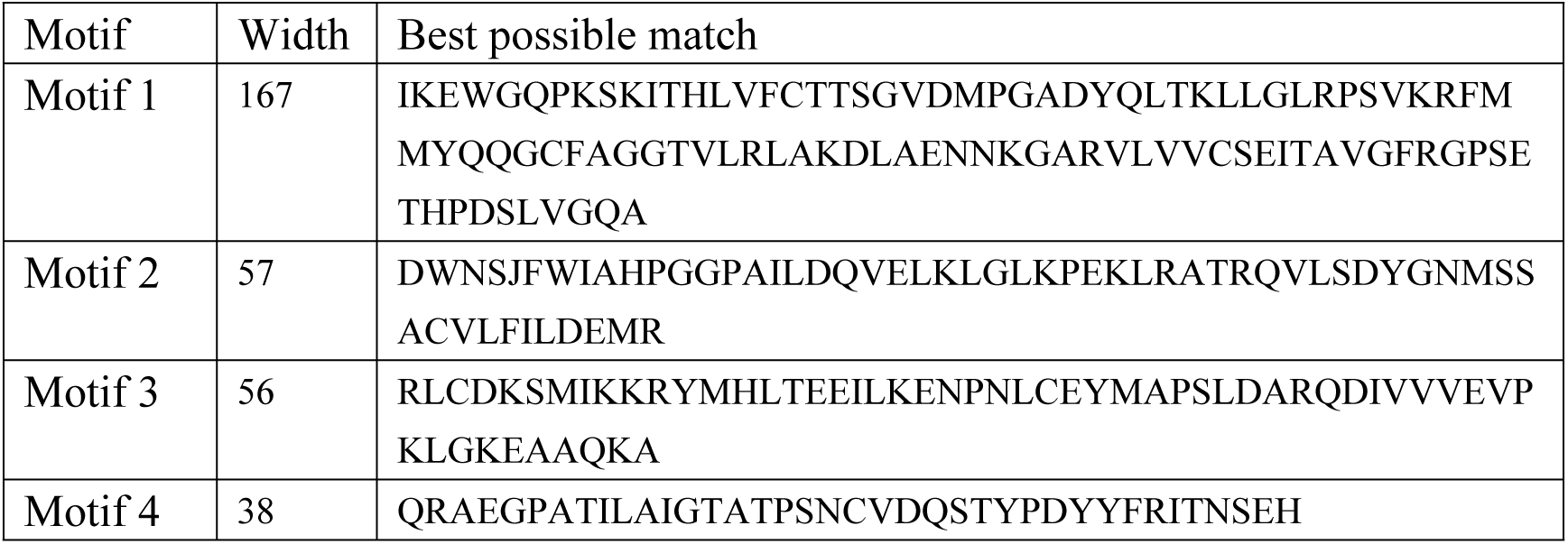
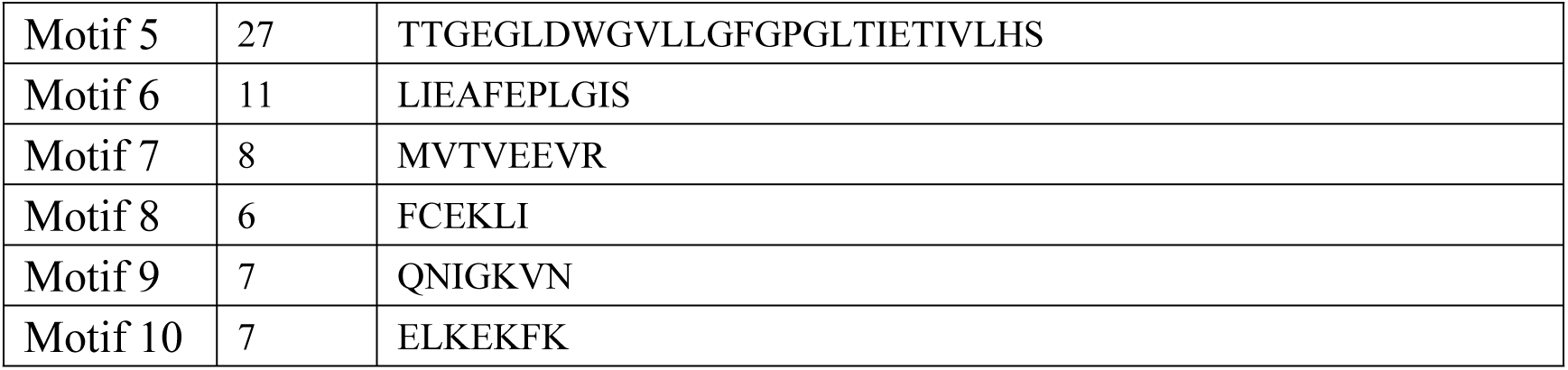
List of the putative motifs of CHS proteins.

**Fig 2.**
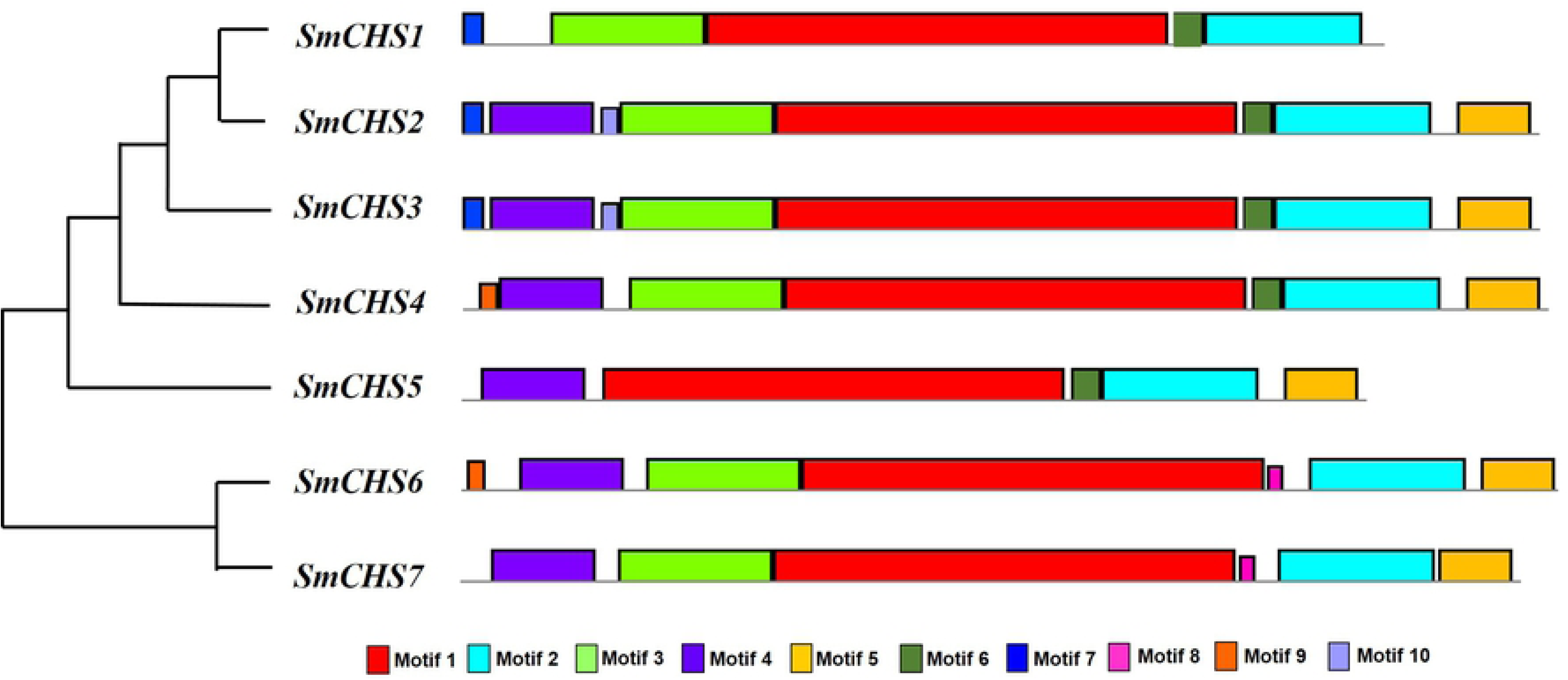
Conserved motifs across all CHS proteins in eggplant. Ten conserved motifs are indicated in different colored boxes.

### Stress-related *cis*-elements in *SmCHS* promoters

To further study the potential regulatory mechanisms of *SmCHS* during abiotic stress responses, the 2 kb upstream sequences from the translation start sites of *SmCHS* were used to identify the cis-elements (Fig 3B). Results showed that all *SmCHS* had common upstream promoter elements, including TATA-box and CAAT-box which occurred more than 100 times, and therefore, these sequences were presumed to be the promoter sequences (Fig 3A). The elicitor response (ERE) and myeloblastosis (MYB) occurred more than 10 times in the *SmCHS* upstream sequences. Research showed that increase of CHS activity causes a high accumulation flavonoid level that inhibit polar auxin transport [15, 38, 39]. Two cis-acting elements (ABRE, involved in abscisic acid responsiveness; AuxRR, involved in auxin responsiveness) were found in the upstream regions. MYB and myelocytomatosis (MYC) binding site have also been identified which may greatly influence plant stress tolerance. These results showed that *SmCHS* is activated by a wide range of environmental and developmental stimuli, and there are many complex ways to regulate *SmCHS* activity in eggplant.

**Fig 3.**
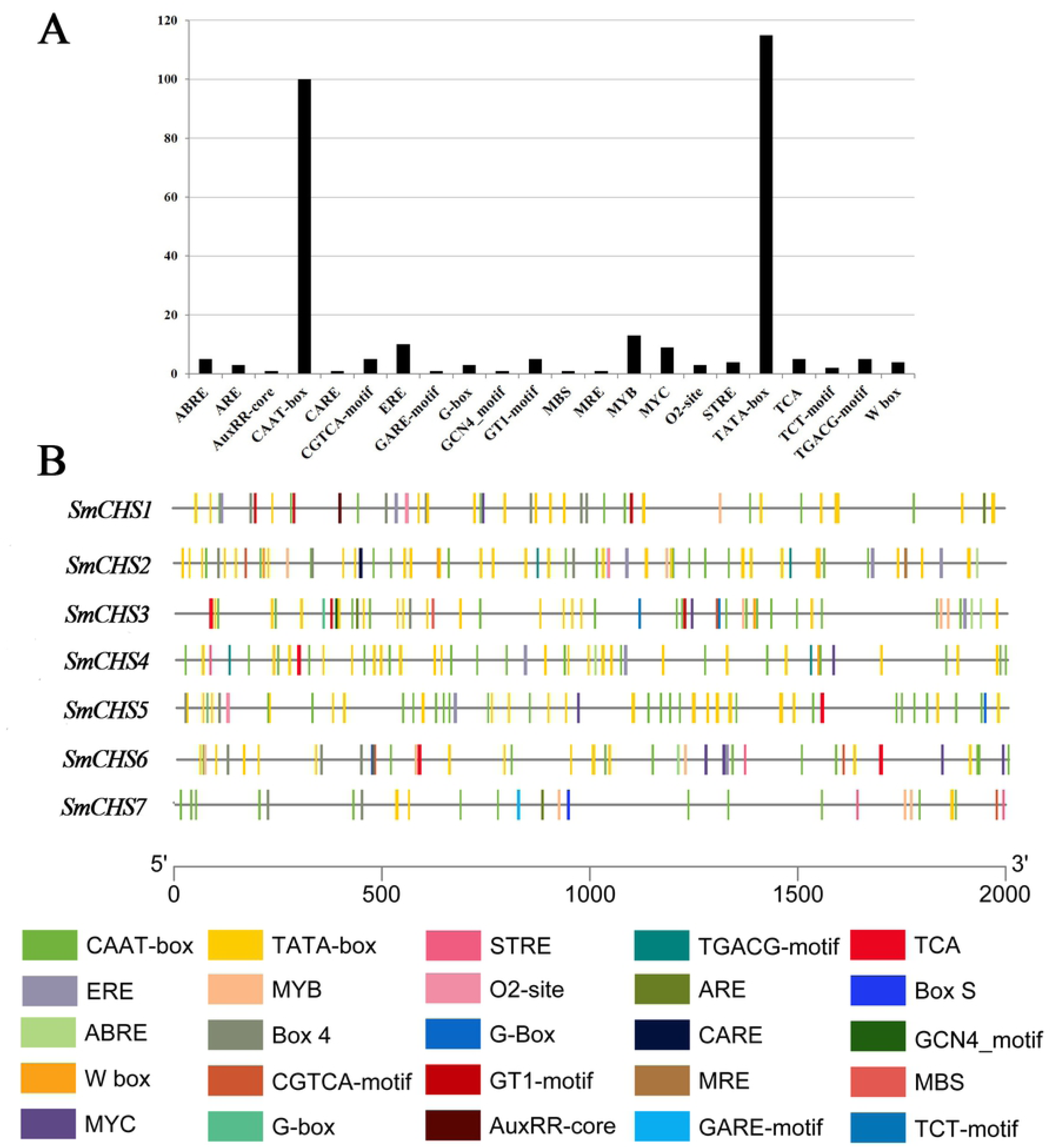
*Cis*-elements in *CHS* family genes promoters. (A) Frequency of cis-element occurrence in upstream sequences. (B) Predicted *cis*-elements in *CHS* genes promoters. Scale bar indicates the length of promoters.

### Phylogenetic analysis of *CHS* genes in Solanaceae

To analyze the evolutionary relationships of CHS genes in Solanaceae, an unrooted phylogenetic tree was constructed using full length amino acid sequences. All the 73 *CHS* genes were classified into 10 groups (Fig 4, Table 3), the number of CHS gene groups ranged from two to eleven. The 7 *SmCHS* genes were categorized into 6 groups (group Ⅰ, Ⅱ, Ⅶ, Ⅷ, Ⅸ and Ⅹ), and the group Ⅱ contained *SmCHS1* and *SmCHS2*. The group Ⅰ, Ⅱ, Ⅸ and Ⅹ exist in all eight species, the group Ⅲ, Ⅳ and Ⅴ were absent in *Solanum melongena* L., *Solanum penellii*, *Solanum lycopersicum* and *Solanum tuberosum* L.. And the group Ⅵ is absent in *Capsicum annuum* L., *Nicotiana tabacum*, *Petunia inflate* and *Petunia axillaries* (Table 3). Those results suggested that the *CHS* genes were conservative, but small variations exist among the eight species in Solanaceae, and also showed that the *SmCHS1*, *SmCHS2* and *SmCHS3* were more conservative than the gene *SmCHS4* according the phylogenetic tree.

**Table 3.** The distribution of *CHS* genes in phylogenetic tree.

| Plant Species | Number | Phylogenetic Group |  |  |  |  |  |  |  |  |  |
| --- | --- | --- | --- | --- | --- | --- | --- | --- | --- | --- | --- |
|  |  | I | II | III | IV | V | VI | VII | VIII | IX | X |
| *Solanum melongena* L. | 7 | 1 | 2 | 0 | 0 | 0 | 0 | 1 | 1 | 1 | 1 |
| *Solanum penellii* | 6 | 1 | 1 | 0 | 0 | 0 | 0 | 1 | 1 | 1 | 1 |
| *Solanum lycopersicum* | 7 | 1 | 1 | 0 | 0 | 0 | 1 | 1 | 1 | 1 | 1 |
| *Solanum tuberosum* L. | 10 | 2 | 1 | 0 | 0 | 0 | 1 | 3 | 1 | 1 | 1 |
| *Capsicum annuum* L. | 9 | 1 | 1 | 0 | 2 | 0 | 0 | 2 | 1 | 1 | 1 |
| *Nicotiana tabacum* | 12 | 2 | 2 | 0 | 0 | 1 | 0 | 3 | 0 | 2 | 2 |
| *Petunia inflata* | 9 | 2 | 1 | 1 | 1 | 1 | 0 | 0 | 1 | 1 | 1 |
| *Petunia axillaris* | 13 | 1 | 1 | 3 | 1 | 3 | 0 | 0 | 1 | 1 | 2 |

**Fig 4.**
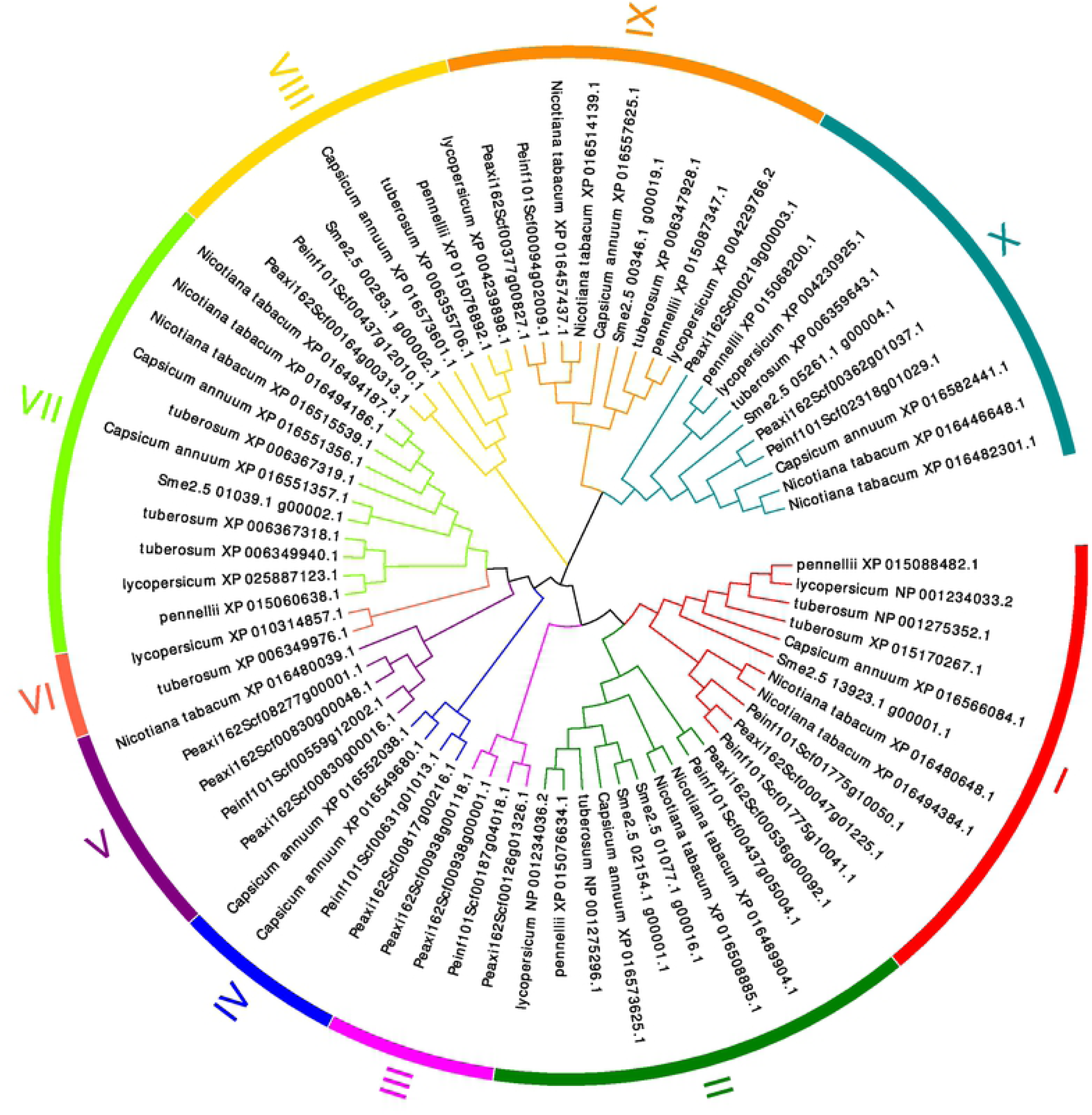
Phylogenetic tree of *CHS* genes in Solanaceae species. The color region is associated with 10 groups of proteins (Group Ⅰ to Ⅹ).

### Expression profile of anthocyanin biosynthesis key genes in eggplant under heat stress

Using the RNA-seq data, a heatmap of 96 anthocyanin biosynthesis key genes (PAL, C4H, 4CL, CHS, CHI, F3H/F3’H, F3’5’H, DFR, ANS, 3GT, MYB1, MYB2, bHLH1, AN11, MADS1) was established under heat stress (Fig 5). The expression of anthocyanidin synthase (ANS) and MYB2 were not identified during this sampling period. For seven *SmCHS* genes, three of them (*SmCHS5*, *SmCHS6*, *SmCHS7*) were not identified, the other four *SmCHS* were divided into two group according their expression patterns. Three of those four *SmCHS* genes (*SmCHS5*, *SmCHS6*, *SmCHS7*) were continuously down-regulated under 38°C and 45°C treatment compared with the CK. However, *SmCHS4* was up-regulated under 38°C but little change at 45°C in peel. And these phenomena have also been observed in some others key gene families associated to anthocyanin biosynthesis.

**Fig 5.**
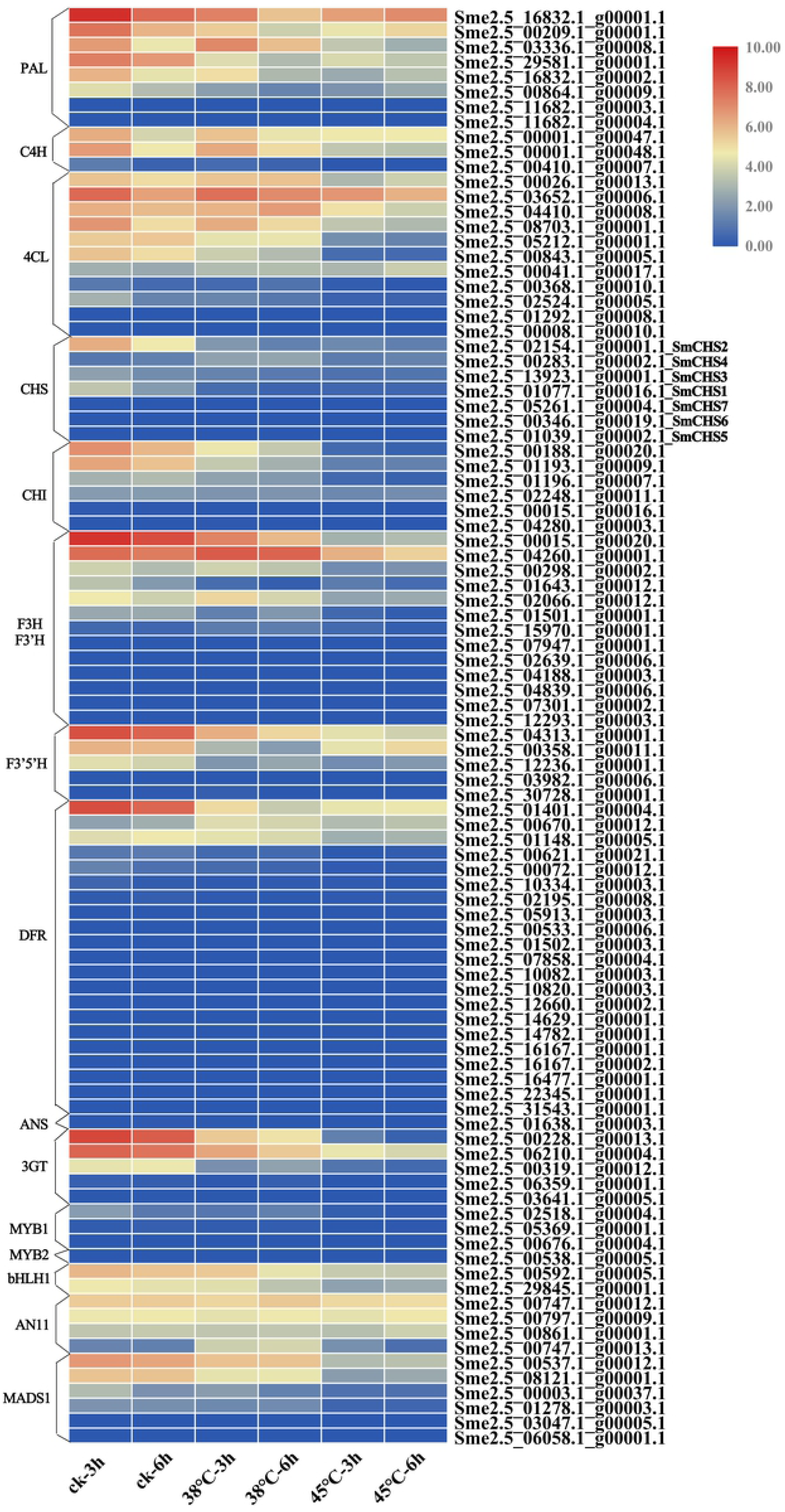
Expression profile of anthocyanin biosynthesis key genes in eggplant under heat stress.

### The mRNA regulatory network associated with anthocyanin biosynthesis in eggplant

A total of 4928 mRNAs correlation coefficient were more than 0.9. all of these mRNAs were functionality categorized in GO database. The top 20 GO enrichment results of biological processes were showed in Table 4. The function was involved in regulation of biological process (GO:0050789), regulation of cellular metabolic process (GO:0031323) and regulation of gene expression (GO:0010468) were collected and filtered to construct regulatory network. Totally, 67 anthocyanin biosynthesis key genes and 146 regulatory mRNAs were included in this regulatory network (S1 Fig). Those GO enrichment results suggest that the anthocyanin biosynthesis pathway maybe regulated by a wide range of environmental and developmental stimuli.

**Table 4.** The top 20 GO enrichment results of biological processes.

| GO term | GO ID | P value |
| --- | --- | --- |
| cellular biosynthetic process | GO:0044249 | 0 |
| cellular nitrogen compound biosynthetic process | GO:0044271 | 0 |
| cellular response to chemical stimulus | GO:0070887 | 0 |
| cellular response to stress | GO:0033554 | 0 |
| regulation of biological process | GO:0050789 | 0 |
| regulation of cellular macromolecule biosynthetic process | GO:2000112 | 0 |
| developmental process | GO:0032502 | 0 |
| regulation of RNA biosynthetic process | GO:2001141 | 0 |
| regulation of cellular metabolic process | GO:0031323 | 0 |
| cellular component organization | GO:0016043 | 0 |
| response to organic substance | GO:0010033 | 1.11E-16 |
| protein metabolic process | GO:0019538 | 1.11E-16 |
| regulation of nitrogen compound metabolic process | GO:0051171 | 1.11E-16 |
| regulation of gene expression | GO:0010468 | 1.11E-16 |
| response to stimulus | GO:0050896 | 1.11E-16 |
| regulation of nucleobase-containing compound metabolic process | GO:0019219 | 1.11E-16 |
| response to chemical | GO:0042221 | 1.11E-16 |
| cell communication | GO:0007154 | 1.11E-16 |
| response to stress | GO:0006950 | 1.11E-16 |
| oxoacid metabolic process | GO:0043436 | 2.22E-16 |

### Expression pattern of anthocyanin biosynthesis key genes in different tissues under heat stress

Using the qRT-PCR data, a heatmap of 20 anthocyanin biosynthesis key genes was established in different tissues under heat stress (Fig 6). The qRT-PCR results showed a high consistency with RNA-seq data, which suggested that RNA-seq data were credible. Most of the CHS genes were expressed in peel, and were low expressed in other tissues. The PAL, 4CL and AN11 genes were mainly expressed in all five tissues. The CHI, F3H, F3’5’H, DFR, 3GT and bHLH1 genes were expressed in flower and peel. The MADS1 were expressed in stem, leaf, flower and peel.

**Fig 6.**
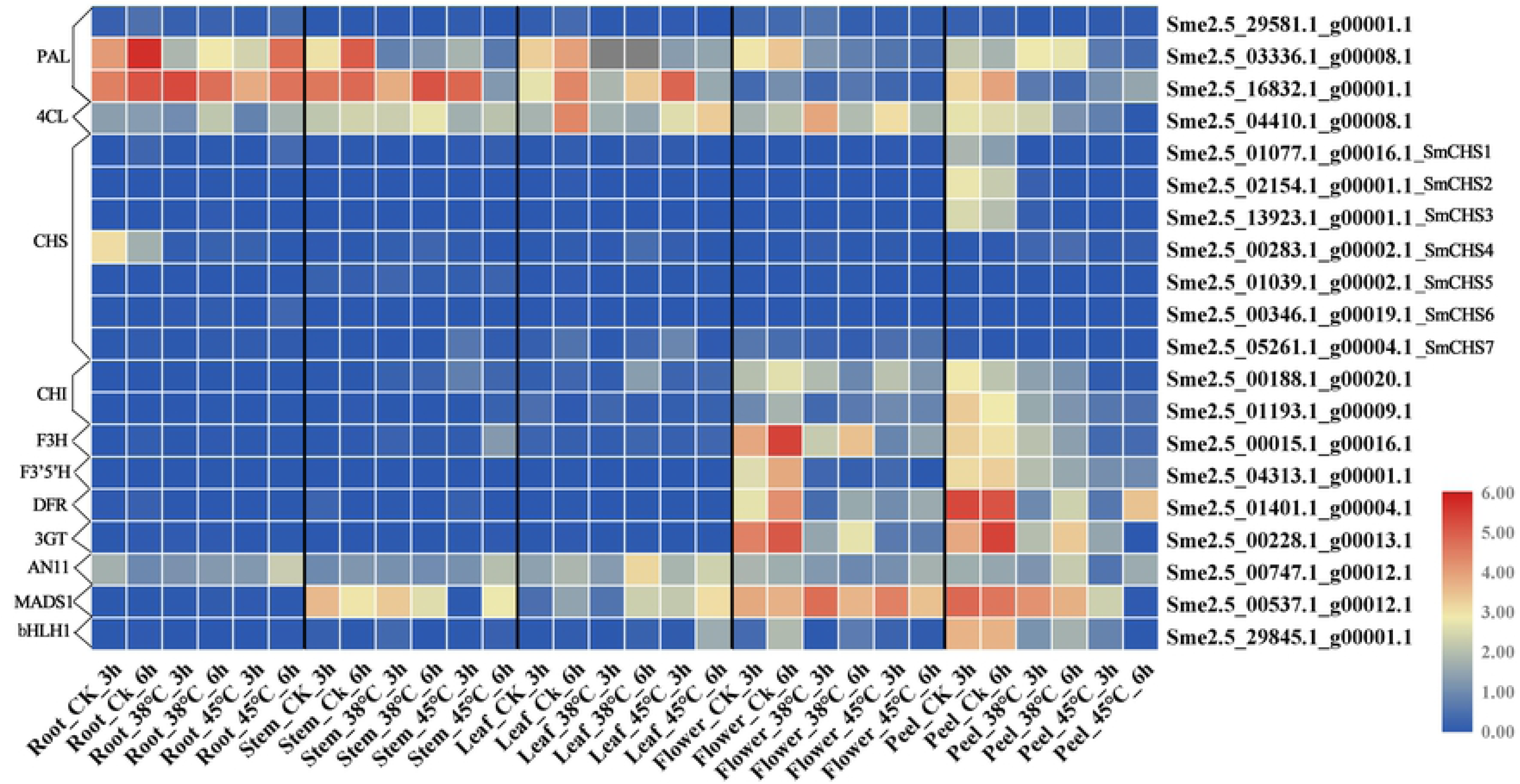
Expression profiles of 20 anthocyanin biosynthesis key genes in different tissues.

### Discussion

It is now known that the *CHS* gene family plays a significant role during the growth and development of plants. In many species, multigene family of *CHS* are identified. For example, Six *CHS* genes have been described in Turnip [40]. In maize, 14 complete *CHS* genes have been identified [41]. Total of 27 *CHS* genes were found in rice [42]. These studies showed that *CHS* members were divided into two or more subclasses according to phylogenetic analysis. Generally, genes grouped into the same subclasses shared similar evolutionary features, and also obtained same expression pattern. In our study, the identified sequences showed the high level of coding sequences similarity (above 90%). The *SmCHS* were classified into three clusters based on the result of maximum-likelihood tree. Under heat stress, cluster i were continuously down-regulation, cluster ii were up-regulated 4 times under 38°C compared with CK in peel, while cluster iii were not detected the expression in eggplant most tissues. Under 35°C, previous studies showed that *SmCHS1* and *SmCHS3* (Sme2.5_01077.1_g00016.1, Sme2.5_13923.1_g00001.1) were down-regulated in peels of eggplant (Lv et al. 2019), which are in agreement with our results. These result suggest that functional diversification of *SmCHS*.

Flavonoids have numerous functions and contribute to pigments, signaling molecules, protectants against biotic and abiotic Stresses. The flavonoid biosynthetic pathway is one of the most intensively investigated pathways for applied biological and genetic processes, as well as for understanding gene regulation, characterization of transposable elements and production of agronomically stress-tolerant plants, and natural dietary antioxidants. biosynthesis of anthocyanins responds to environmental stressors such as light, nutrient depletion, and temperature change. The peel color determined by the content of anthocyanin, is a majority economic trait for eggplant, which are modulated by the genes in flavonoid biosynthesis pathway. Compared with the other tissues, *SmMYB1* and all anthocyanin biosynthetic key genes (*SmCHS*, *SmCHI*, *SmF3H*, *SmDFR*) except *SmPAL* were dramatically upregulated in the fruit skin of the purple cultivar [43]. The full lengh cDNA of *SmCHS*, *SmCHI*, *SmF3’5’H*, *SmDFR*, were isolated from eggplant by Jiang. These genes have highest expression levels in peels except for *SmF3H* which was detected in stems [44]. The expression profiles of these key gene families under heat stress were investigated in our study. The PAL, 4CL and AN11 genes were mainly expressed in all five tissues. The CHI, F3H, F3’5’H, DFR, 3GT and bHLH1 genes were expressed in flower and peel, inferred that these genes respond at late-stage of anthocyanin pathway directly regulate the color of fruit skin and flower.

Heat stress reduced the anthocyanin content and the enzyme activities of CHS, DFR, ANS and 3GT/UFGT in eggplant peel, and strengthened the activity of PAL [34, 45]. When the temperature is over 35°C, the eggplant will be dehydrated and shrink, and peel’s color will turn light. CHS is a key enzyme of the flavonoid biosynthesis pathway. Most of the genes associated with flavonoid biosynthesis were down-regulated under heat stress. In this study, the genes of the flavonoid biosynthesis pathway showed tissue-specific and genes express in different phases and tend to change with the time (Fig 6). According to the RNA-seq results of 96 anthocyanin biosynthesis key genes in eggplant peel, *SmCHS4* showed the highest expression level at the 38°C-3h along with eight other genes (*Sme2.5_03336.1_g00008.1_PAL*, *Sme2.5_00041.1_g00017.1_4CL*, *Sme2.5_00283.1_g00002.1_smCHS4*, *Sme2.5_00298.1_g00002.1_F3H*, *Sme2.5_02066.1_g00012.1_F3H*, *Sme2.5_04260.1_g00001.1_F3H*, *Sme2.5_15970.1_g00001.1_F3H*, *Sme2.5_00670.1_g00012.1_DFR*, *Sme2.5_00747.1_g00013.1_AN11*) (Fig 7). Particularly, *Sme2.5_03336.1_g00008.1_PAL* expression level under 38°Cdoubled while down-regulated in 45°C compared with CK; *Sme2.5_00670.1_g00012.1_DFR*, *Sme2.5_00747.1_g00013.1_AN11* expression level increased 3-4 fold and 7-10 fold under 38°C, respectively. In addition, 52 genes expression level were reduced under heat stress which were similar to Lv’s results [9], while 35 genes expression were not identified. Those results suggested that some anthocyanin biosynthesis key genes contribute to protect the eggplant from the damage of heat stress. Moreover, these genes families exhibited two or more expression patterns and execute multiple genetic functions to regulate anthocyanin content. Combined with regulatory networks, it is possible to further understand the regulation mechanism of peel color in eggplant.

**Fig 7.**
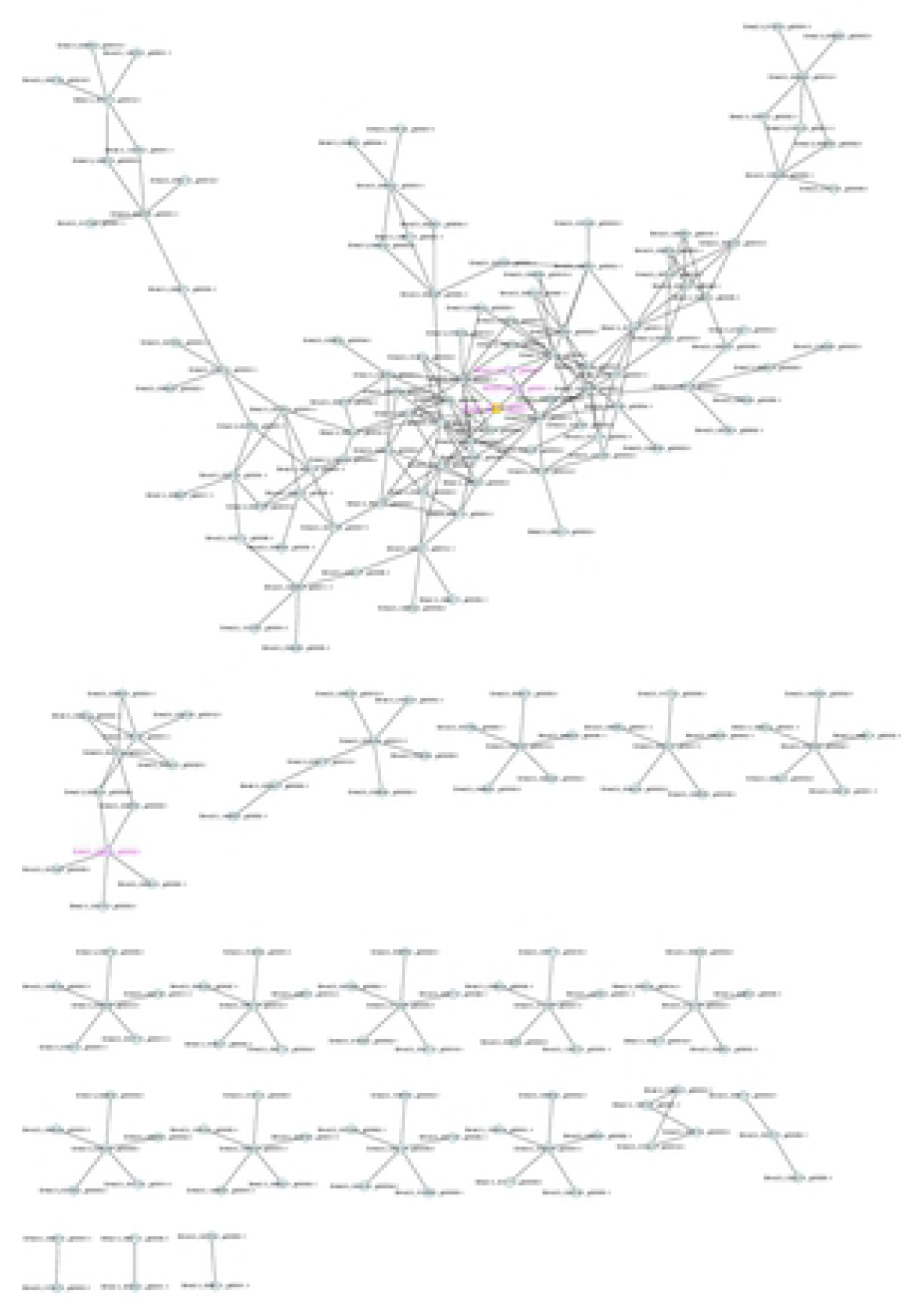
The anthocyanin biosynthesis key gene expression profiles in response to heat stress which were observed the highest expression level at 38°C-3h.

## Conclusions

In this study, a genome-wide analysis about *SmCHS* genes family in eggplant was performed. The CHS protein biochemical characteristics, phylogenetic relationships, gene structures, *cis*-regulatory elements, regulatory network and functional predictions of the family members were examined. It is revealed that *SmCHS* genes family have conservation gene structure and functional diversification. CHS play important roles to anthocyanin biosynthesis pathway, exhibited two or more expression patterns and execute multiple functions to regulate anthocyanin content in eggplant peel under heat stress. This work will make eggplant for further research of functions, regulation and evolution of the CHS family.

## Acknowledgments

This work was supported by Agricultural Committee Basic Project (Shanghai Agricultural word (2015) No 6–2-3), the National Key Technology R&D Program during the 13th Five-Year Plan Period (2017YFD0101904) and China Agriculture Research System (Grant No. CARS-25). The funding bodies didn’t play a role in the design of the study and collection, analysis, and interpretation of data and in writing the manuscript.

## Author contribution

DZ put forward the research, ZZ and AD carried out the preparation and treatment of test materials. XL and SJ designed and performed the experiment. XW and SZ analyzed the data and wrote the manuscript. XW revised the article. All authors read and approved the final manuscript.

## Supporting Information

**S1 Table**. The CHS protein sequences of Solanum species.

**S2 Table**. Features of CHS genes identified in Solanum species.

**S3 Table**. Primers used for real time PCR analysis.

**S1 Fig**. The interaction network key with anthocyanin biosynthesis in eggplant. The pink labels represent the CHS gene family.

---

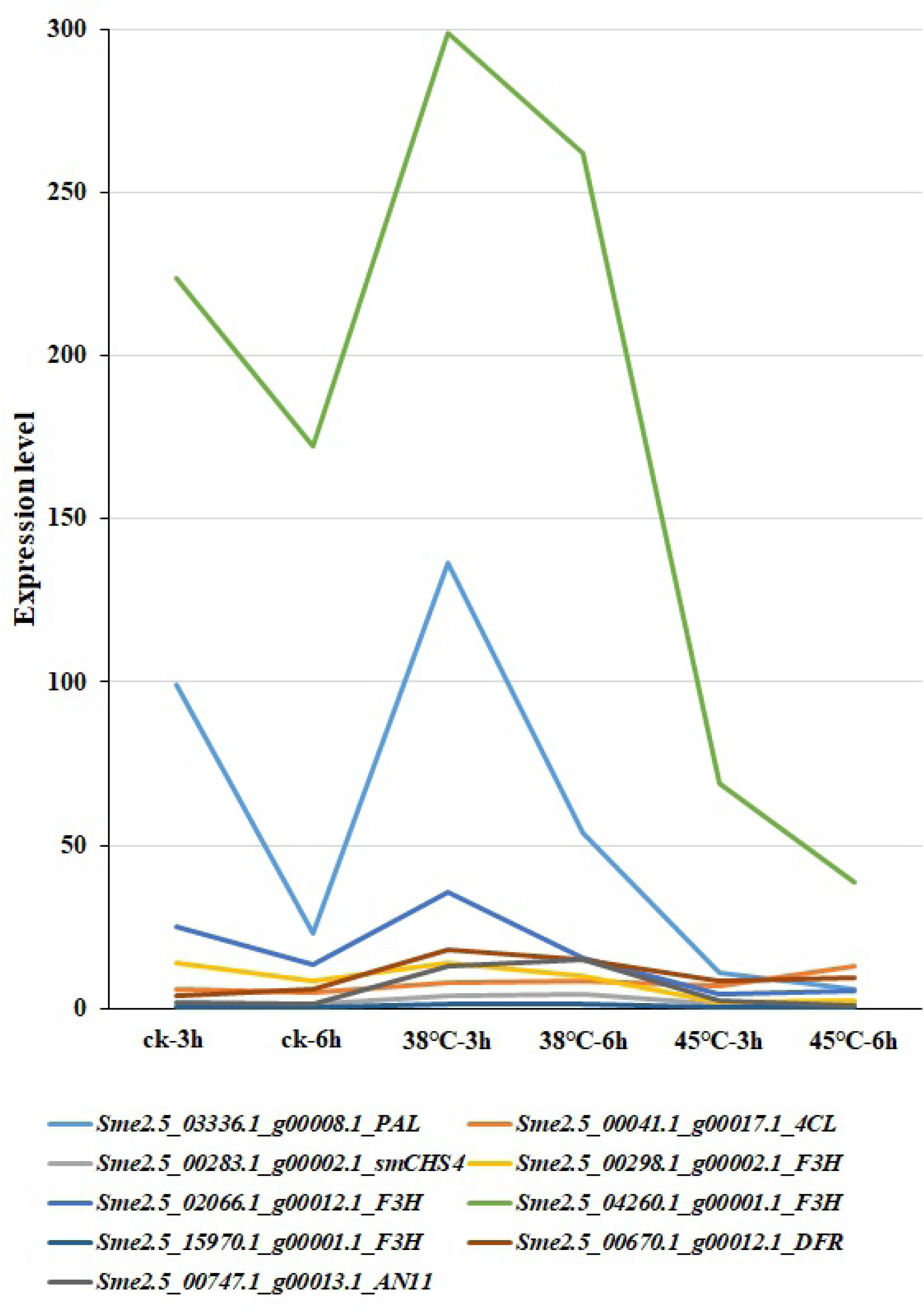

